# How many are you? Open data and bioinformatics reveal species misidentification and potential introgression in *Chordodes* (Phylum Nematomorpha)

**DOI:** 10.64898/2026.02.03.703548

**Authors:** Mattia De Vivo

## Abstract

The potential usage of genomic open data can help us to understand patterns in biodiversity. They can also be helpful for identifying morphologically similar species. An example of taxon in which this can be useful is Nematomorpha, one of the less studied animal phyla, for which data has started to be available recently and where species identification can be hard. In this study, I planned initially to evaluate the usage of mitochondrial data for population analyses using an RNA sequencing (RNA-seq) dataset labelled as belonging to *Chordodes fukuii*. After surprising results using extracted sequences from the barcoding gene cytochrome c oxidase subunit I (COXI), I evaluated species delimitation using a mix of a previously released double-digest restriction-site-associated DNA sequencing (ddRADseq) SRA dataset plus the RNA-seq one. PCA, R analyses through “adegenet” and ADMIXTURE confirmed the presence of two species in the RNA-seq dataset, which should be labelled as *C. formosanus* and *C. japonensis*; however, some individuals labelled as *C. japonensis* according to COXI clustered with *C. formosanus*’s specimens or had some *C. formosanus’* ancestry when more data was used, indicating potential introgression or incomplete lineage sorting. The study shows how previously released data can be used for evaluating species delimitation, potential previous demographic events and potential needs in DNA barcoding and genomics for avoiding future misidentification of morphologically similar species.

## Introduction

Recent improvements in genomic sequencing led to the accumulation of data in open repositories, which in turn can be used for analyses and infer new patterns (Plocik & Graveley, 2013). Such data can be used to understand biodiversity or help in exploring it. For example, data from open repositories have been used to assemble whole mitogenomes for inferring species relationships (e.g., Vieira & Prosdocimi 2019), analyze potential genomes rearrangements (e.g., Lewin *et al*. 2023) and infer potential gene loss in lineages, e.g. parasites (e.g., Herlyn *et al*. 2025).

Among organisms that have seen an increase of available data thanks to the sequencing improvement, horsehair worms (phylum Nematomorpha) are slowly getting more uploaded DNA and RNA sequences and raw data available after decades of almost complete neglect. Specifically, using GenBank (last accessed: June 24, 2025), one of the most popular data repositories for nucleotide data (Clarke *et al*., 2016) as reference, 11 genomes are available; an increased number compared to the absolute zero that was present before 2023, when three sequenced genomes were made available (Cunha *et al*., 2023; Eleftheriadi *et al*., 2024). In addition, Sequencing Read Archives (SRAs) from several projects have been uploaded, for a total of 16 BioProjects with at least a hairworm species sequenced and 103 BioSamples available for species inside this phylum. So far, the released data showed how horsehair worms lost ortholog metazoan genes (Cunha *et al*., 2023; Eleftheriadi *et al*., 2024), which seem to be common in other parasitic Metazoa (Herlyn *et al*. 2025), and how their mitogenomes have inverted repeats in some protein-coding genes, which are unusual for animals (Mikhailov *et al*., 2019). In addition, transcriptomic data has been showing potential horizontal gene transfers between freshwater hairworms (class Gordioida) and their hosts, which probably helps the parasites to “manipulate” their hosts (Mishina *et al*., 2023).

The availability of nucleotide data could also help us to understand horsehair worms’ ecology and taxonomy better. Specifically, although they have a fearsome reputation as “body snatchers” (albeit not all the taxa induce their hosts to jump into water; see Schmidt-Rhaesa 2013 and references therein), Nematomorpha’s popular science notoriety does not translate into a research interest and the phylum is widely regarded as one of the most understudied animal groups (Schmidt-Rhaesa, 2013). Given this, we have a relatively limited grasp of their ecology and systematic relationships. For example, the hosts are not known for some species (Schmidt-Rhaesa, 2013 and references therein). In addition, horsehair worms have very few morphological characters for taxon identification; because of this, relationships among genera or species in the same genus are very often unclear (Schmidt-Rhaesa, 2013; Bolek & Hanelt, 2024). However, when molecular data were available, it was shown how species complexes or even “genera complexes” are present (Anaya *et al*., 2021; Bolek & Hanelt, 2024; Groß *et al*., 2025). Therefore, having genomic data can help us understand their phylogeny and if such phylogenetic relationships have also an ecological meaning: for example, Bolek and Hanelt (2024) showed how species in the *Gordius robustus* species complex tend to phylogenetically cluster together according to parasitized final host, stressing the potential use of phylogeny for uncovering ecological interactions.

In this paper, I show how to use a dataset from a previous publication for getting barcodes useful for identifying species and try to confirm the species ID by combining more data together. For doing so, I extracted cytochrome c oxidase subunit I (COXI) sequences from the SRAs of Mishina *et al*. (2023); such SRAs were labelled as *Chordodes fukuii*, which is one of the horsehair worms present in Japan (Inoue, 1952; Schmidt-Rhaesa, 2004) and I wanted to understand their population dynamics through mitochondrial data together with their host, in a similar fashion as De Vivo *et al*. (2023a). After using BLAST (Altschul *et al*., 1990), I got surprisingly high (≈100%) BLAST percentage IDs for two different species (Sup. Table 1). I then adapted the RNA data together with a double-digest restriction-site-associated DNA sequencing (ddRADseq) dataset generated by another study (De Vivo *et al*., 2023a). By using this dataset, I tried to confirm the original COXI identification and check potential demographic events. Notice that, although here the analyses are tailored for a specific horsehair worm group, the pipeline could virtually be used for any species of any life kingdom.

## Materials and Methods

### Used HPC environment and software installation

The workflow followed in this study is summarized in Fig. 1 and its codes are available in Sup. Data.

**Figure 1.**
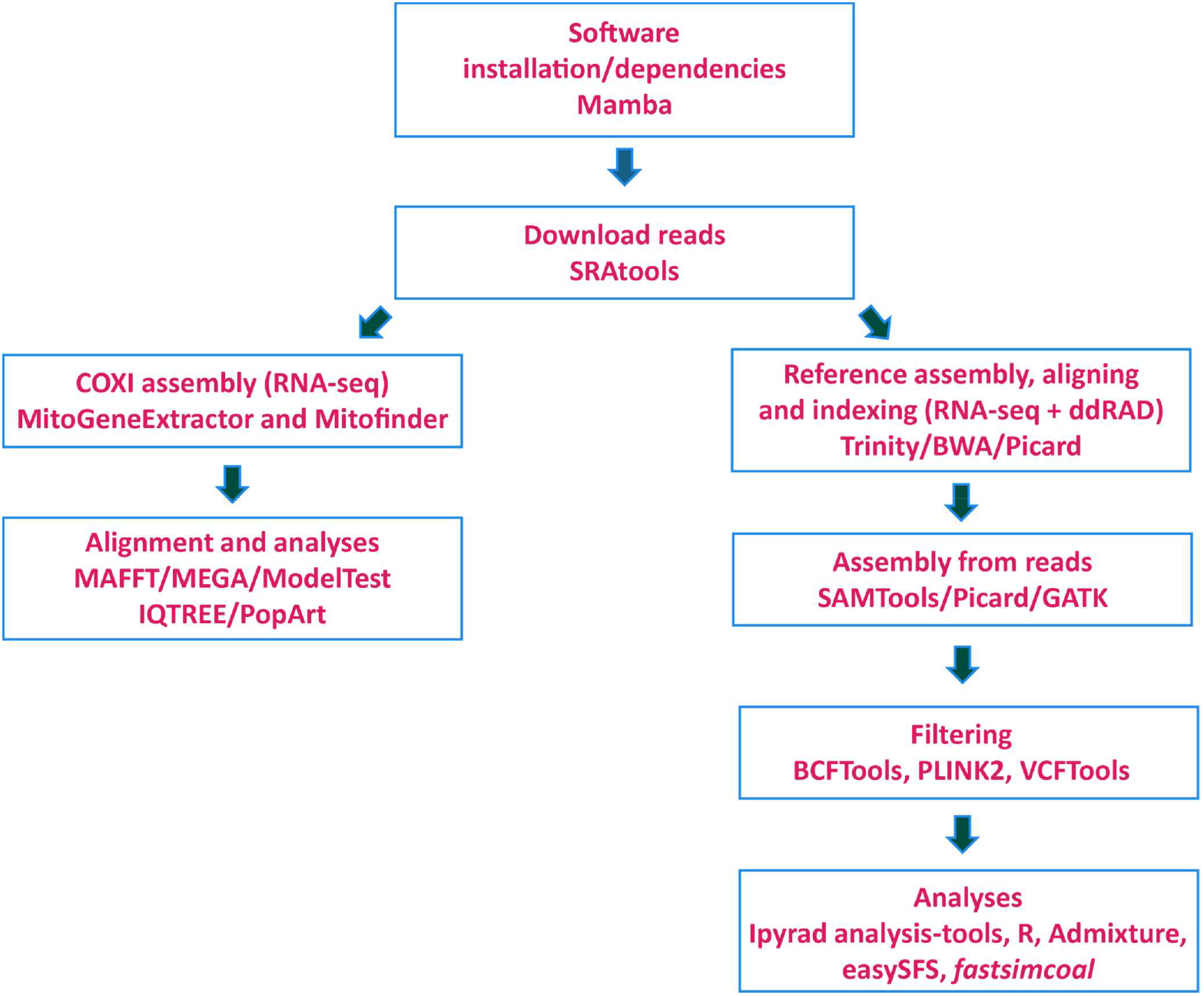
Summary of the bioinformatic workflow followed in this paper.

The reads’ download and their assemblies were done by using the High Performance Computing (HPC) at the University of Kaiserslautern-Landau (https://hpc.rz.rptu.de/) in a SLURM environment. I installed the used software and/or the needed dependencies in different Conda environments in the HPC by using Mamba (version 1.5.9; https://github.com/mamba-org/mamba).

### RNA dataset and initial COXI assembly

Using the SRA Toolkit (version 3.1.1; https://github.com/ncbi/sra-tools), I downloaded the *Chordodes* RNA-seq data from Mishina *et al*. (2023) (BioProject: PRJNA994131). Given that the RNA-seq data was already trimmed of low-quality reads and adapter sequences, I did not perform any additional trimming.

For assembling protein-coding mitochondrial genes, I used MitoGeneExtractor (version 1.9.5; Brasseur *et al*., 2023) using the protein annotation of the only *Chordodes* mitogenome available in GenBank at the time of the analysis (accession number: MG257764) as a reference. Given that the SRAs were trimmed, it was not possible to recover all the genes from them (see Brasseur *et al*. 2023): therefore, I decided to keep only the COXI data for further analyses, given that most of the mitochondrial data of horsehair worms in GenBank belongs to this gene. For two specimens that presented some “N” in their COXI assembly (SRR25249029 and SRR25249043) and for an additional specimen that did not cluster in the analyses with more genes as it did with COXI (SRR25249039), an additional assembly was attempted by using MitoFinder (version 1.43, Singularity version; Allio *et al*., 2020) using the GenBank (gb) version of the same mitogenome used for MitoGeneExtractor as reference.

### BLASting of the COXI sequences, phylogenetic tree and haplotype network

For checking if I was able to get true *Chordodes* sequences from the reads, I used the online version of BLAST (Altschul *et al*., 1990), with the BLASTn algorithm. After getting 100% IDs on sequences in the database (Sup. Table 1), I downloaded all the available sequences from such species (Sup. Table 2). I also downloaded sequences from an unidentified *Chordodes* species (Kuroda *et al*., 2024; Sup. Table 2).

After downloading such sequences, I aligned them together with the COXI from the used reference mitogenome and the ones generated from the RNA-seq data using MAFFT (version 7.526; Katoh & Standley, 2013) with the L-INS-i algorithm. I manually trimmed such sequences while visualizing them with MEGA11 (version 11.0.13; Tamura *et al*., 2021). For building a phylogenetic tree, I first checked the most likely nucleotide substitution model using ModelTest-NG (version 0.1.7; Darriba *et al*., 2020) implemented in raxmlGUI (version 2.0.13; Edler *et al*., 2021). I used the most likely model (JC69; Jukes & Cantor, 1969) according to the small-sample corrected Akaike Information Criterion (AICc; Sugiura, 1978; Sup. Data) for building a phylogeny using IQ-TREE (version 2.2.6; Minh *et al*., 2020) with 1000 ultrafast bootstrap replicates (UFBoot; Hoang *et al*., 2018). I visualized the tree with FigTree (version 1.4.4; Rambaut, 2018) and rooted it with midpoint rooting.

For understanding and visualizing the diversity of the sequences better, I generated a haplotype network with the TCS algorithm (version 1.21; Clement *et al*., 2000) with PopArt (version 1.7; Leigh & Bryant, 2015) by using the same alignment used for the phylogenetic tree.

### Assembling of the RNA and ddRAD data and VCF filtering

After getting the phylogenetic tree, I downloaded the 27 ddRAD sequencing data used in the final dataset for *Chordodes formosanus* by De Vivo *et al*. (2023a; Sup. Table 3). This data was uploaded already trimmed too, so no trimming was necessary. I then assembled the RNA-seq and ddRAD reads following a modification of the pipeline followed in Fuhrmann *et al*. (2023). First, I assembled the transcriptome of the individual SRR25249029, which was the one with most reads from the “after” class (i.e., the worm was extracted from the host and then left for a week in the water; see Mishina *et al*. 2023 and the GEO dataset of the paper) by using Trinity (version 2.15.2; Haas *et al*., 2013). Such transcriptome was indexed, and the downloaded reads were aligned to it; both processes were done through BWA (version 0.7.18; Li & Durbin, 2010). A following dictionary file of the transcriptome was created with picard (version 3.3.0; https://broadinstitute.github.io/picard).

The SAM files resulting from the alignment were converted into BAM, sorted and indexed by SAMTools (version 1.18; Danecek *et al*., 2021). The sorted BAM files were initially processed through picard with the “FixMateInformation”, “MarkDuplicatesWithMateCigar” and “AddOrReplaceReadGroups” commands and then indexed through SAMTools. The genotypes were called and outputted in a Variant Call Format (VCF) file through GATK (version 3.8; Van der Auwera *et al*., 2013), using the “HaplotypeCaller”, “BaseRecalibrator”, “PrintReads”, “CombineGVCFs” and “GenotypeGVCFs” functions. I only kept individuals with at least 5000 retained reads in the final VCF (Sup. Data), which led me to have 27 individuals: 8 from ddRAD and 19 from RNA-seq (Table 1).

**Table 1.** Individuals included in the final GATK VCF assembly.

| SRA ID | Source | Type of data |
| --- | --- | --- |
| SRR22822833 | De Vivo et al. (2023a) | ddRAD |
| SRR22822835 | De Vivo et al. (2023a) | ddRAD |
| SRR22822837 | De Vivo et al. (2023a) | ddRAD |
| SRR22822842 | De Vivo et al. (2023a) | ddRAD |
| SRR22822849 | De Vivo et al. (2023a) | ddRAD |
| SRR22822852 | De Vivo et al. (2023a) | ddRAD |
| SRR22822860 | De Vivo et al. (2023a) | ddRAD |
| SRR22822862 | De Vivo et al. (2023a) | ddRAD |
| SRR25249025 | Mishina *et al.* (2023) | RNA-seq |
| SRR25249026 | Mishina *et al.* (2023) | RNA-seq |
| SRR25249027 | Mishina *et al.* (2023) | RNA-seq |
| SRR25249028 | Mishina *et al.* (2023) | RNA-seq |
| SRR25249029 | Mishina *et al.* (2023) | RNA-seq |
| SRR25249030 | Mishina *et al.* (2023) | RNA-seq |
| SRR25249031 | Mishina *et al.* (2023) | RNA-seq |
| SRR25249032 | Mishina *et al.* (2023) | RNA-seq |
| SRR25249033 | Mishina *et al.* (2023) | RNA-seq |
| SRR25249034 | Mishina *et al.* (2023) | RNA-seq |
| SRR25249035 | Mishina *et al.* (2023) | RNA-seq |
| SRR25249036 | Mishina *et al.* (2023) | RNA-seq |
| SRR25249037 | Mishina *et al.* (2023) | RNA-seq |
| SRR25249038 | Mishina *et al.* (2023) | RNA-seq |
| SRR25249039 | Mishina *et al.* (2023) | RNA-seq |
| SRR25249040 | Mishina *et al.* (2023) | RNA-seq |
| SRR25249041 | Mishina *et al.* (2023) | RNA-seq |
| SRR25249042 | Mishina *et al.* (2023) | RNA-seq |
| SRR25249043 | Mishina *et al.* (2023) | RNA-seq |

I removed indels and filtered for minor allele frequency (MAF) and maximum alleles from the VCF through the “view” function of BCFTools (version 1.21; Danecek *et al*., 2021), setting MAF at 0.02 and maximum alleles at 2. I then pruned the VCF for linkage disequilibrium (LD) with PLINK2 (version 2.0.0a.6.9; www.cog-genomics.org/plink/2.0/), with the “indep-pairwise” function, using a 50,000 base pairs (bps) window in which LD was checked every 10,000 bps after each step, with a variance inflation factor (VIF) set at 0.5 (“indep-pairwise 50 10 0.5”).

Finally, I removed loci with more than 25% missing data from the VCF with VCFTools (version 0.1.16; Danecek *et al*., 2011), with the “max-missing” function set at 0.75 and kept only variable sites with the “recode” function.

### Population structures’ analyses with the VCF file

For analyzing the final datasets and understanding the number of clusters (K) in the VCF, I employed three strategies. The first one used ipyrad (version 0.9.105; Eaton & Overcast, 2020) and its analysis-tools suite (https://ipyrad.readthedocs.io/en/latest/API-analysis/index.html). First, the VCF file was converted into a HDF5 database, with the “ld_block_size” argument set as 50,000 (https://ipyrad.readthedocs.io/en/latest/API-analysis/cookbook-vcf2hdf5.html). Then, I ran a *k-means* principal component analyses (PCA; https://ipyrad.readthedocs.io/en/latest/API-analysis/cookbook-pca.html), with 100 replicates, 3 populations defined in the “imap” dictionary (Sup. Data), the “minmap” parameter set at 0.5 and the “mincov” parameter set at 0.7.

As a second approach, I used functions inside R (version 4.3.1; R Core Team, 2023) implemented in RStudio (version 2023.06.1; Posit Team, 2023). I used the package “vcfR” (version 1.15.0; Knaus & Grünwald, 2017) for loading the VCF in R and converting it to the genind format. I then used the snapclust clustering method and the “find.clusters” function implemented into “adegenet” (version 2.1.10; Jombart *et al*., 2010; Beugin *et al*., 2018) for evaluating the ideal number of clusters. In the case of the latter, I used all the available components, and I used the generated data for bar plots. Finally, I modified the chromosome information in the VCF through BASH scripts for making it compatible with ADMIXTURE (32-bit version, version 1.3.0; Alexander *et al*., 2009; Sup. Data.) and I ran the software, following the scripts available in the manual (https://dalexander.github.io/admixture/admixture-manual.pdf) for Ks ranging from 1 to 3, choosing the best K according to crossvalidation. All the bar plots (both with data from the “find.clusters” function and the ones from ADMIXTURE) were done in R through the “xadmix” package (version 1.0.0; Schönmann, 2022).

After noticing some individuals that clustered with a different group compared to COXI in the ipyrad PCA-”find.clusters” comparison and that had roughly 80:20 assignment to clusters when K=2 with ADMIXTURE, I checked the number of missing sites per individual with the “missing.sort_values” function of the ipyrad analysis-tools, since it is known missing data can affect population assignment (e.g., Yi & Latch, 2022). I then re-tried to run the three previously described approaches with a VCF without those individuals that had exactly or more than 25% missing data (i.e., SRR25249032, SRR25249035 and SRR25249039; Sup. Table 4), which were removed in R through “vcfR” using scripts from Dalapicolla *et al*. (2021; available at https://github.com/jdalapicolla/LanGen_pipeline_version2/blob/master/PART1/1.1_FILTERING_SNPs.R).

### Demographic events

For understanding if there was a hybridization event in the past, I used *fastsimcoal* (version 2.8.0.0; Excoffier *et al*., 2021) using the same models made by Meier *et al*. (2017; see https://speciationgenomics.github.io/fastsimcoal2). Briefly, these models were: “Early gene flow” (i.e., there was hybridization in the past and then it stopped soon after); “No geneflow”; “Recent gene flow” (gene flow in recent times due to secondary contact or similar); “Different gene flow” (different rates of gene flow between old and recent times); “Constant (i.e., ongoing) gene flow”. Compared to the original models, I substituted the 6,000 generations with a potential first hybridization event (TDIV) with a uniform distribution between 1 and 100,000; “1” is the generation time of *Chordodes* (1 year; Chiu, 2017), while 100,000 is a time set after checking a population drop according to Stairway Plots in De Vivo *et al*. (2023a), which may correspond to a population split. However, I preferred to not set that parameter as bounded for allowing potential estimates outside of such a boundary. In addition, it is highly likely that the original hybridization event is far more ancient, given the old age of Nematomorpha (Poinar & Buckley, 2006; Howard et al., 2022). Per each model, I set the mutation rate as 2×10^9^ (see De Vivo *et al*. 2023a for more details).

For running the software, I followed the same methodology as De Vivo *et al*. (2023b); briefly, I generated two SFS files using easySFS (version 0.0.1; https://github.com/isaacovercast/easySFS): one with all the outputted individuals and one with the samples that had less than 25% missing data. Therefore, I had 2 SFS files per each model. Although easySFS takes into consideration missing data, I preferred to see if excluding individuals would have changed the models’ results. In both cases, I selected all the potential *C. japonensis* individuals (defined by COXI, i.e., CJPOT: see the figures in the manuscript), while I chose the number of *C. formosanus* individuals that maximized the number of segregating sites (see the repository’s website for more detail). After this, I run *fastsimcoal* with 100 replicates per each model, with 200,000 coalescent simulations (-n) and 50 optimisation cycles (ECM; -L) for estimating parameters based on maximum likelihood (-M). The analyses considered the monomorphic sites by pooling together SFS entries with less than 10 SNPs (-C 10) and outputting all SNPs (-s0).

After the models were run, I chose the run with the best estimate for each model with a bash script (available at https://raw.githubusercontent.com/speciationgenomics/scripts/master/fsc-selectbestrun.sh). From such runs, I calculated the Akaike information criterion (AIC; Akaike, 1998) using the highest likelihood values from each model by converting the log10-likelihoods reported by *fastsimcoal2* to ln-likelihoods, using the modified script from De Vivo *et al*. (2023b). I finally calculated AIC weights (Wagenmakers & Farrell, 2004) among the two model pairs using the “akaike.weight()” function of the R package ‘qpcR’ (version 1.4-1; Ritz & Spiess, 2008). I also checked the likelihood distributions for model comparison. From the best model of each SFS, I then run 100 parameter estimation analyses by using 100 bootstrapped VCF files, following scripts from De Vivo *et al*. (2023b). I then took the values of the time of the events according to the most likely model and made a confidence interval based on mean and standard deviation (SD) through the R “base” package (version 4.3.1; R Core Team, 2023). I made box plots with the “tidyverse” package (version 2.0.0; Wickham *et al*., 2019) for visualizing the difference in estimated values according to the replicates.

In addition to *fastsimcoal*, I used triangle plots for evaluating the presence of hybrids and backcrosses in my samples with the R package “triangulaR” (version 0.0.1; Wiens *et al*., 2025). For doing this, I used the vcfR version of the generated VCF files used also for ADMIXTURE and I evaluated the results with “1” as difference threshold.

## Results

### COXI results

In total, 200 different sequences were used for the COXI analyses, with 352 bps each. The phylogenetic tree shows how the sequences extracted from the RNA-seq dataset do not cluster with each other, forming two groups with sequences belonging to *Chordodes formosanus* and *C. japonensis* (bootstrap support=100; Fig. 2A; Sup. Data.). The top BLAST results show that the sequences differ too: each top result matches the IDs from the tree (Sup. Table 1). The haplotype network also shows this pattern, since the sequences do not cluster together and instead form haplotype groups with previously released *C. formosanus* and *C. japonensis* sequences (Fig. 2B). The BLAST results, the output of ModelTest-NG and the haplotype network show how eight COXIs extracted from RNA-seq data (SRR25249025, SRR25249026, SRR25249028, SRR25249030, SRR25249034, SRR25249036, SRR25249037 and SRR25249038) are 100% identical to previously released Japanese *C. formosanus* sequences, while 6 sequences (SRR25249029, SRR25249031, SRR25249032, SRR25249033, SRR25249035 and SRR25249039) are 100% identical to publicly available *C. japonensis* sequences (Sup. Table 1; Sup. Data). 5 other extracted sequences (SRR25249027, SRR25249040, SRR25249041, SRR25249042 and SRR25249043) were almost 100% equal to previously released *C. japonensis* sequences (BLASTn similarity percentage ranging between 99.68 and 99.848%; Sup. Table 1).

**Figure 2.**
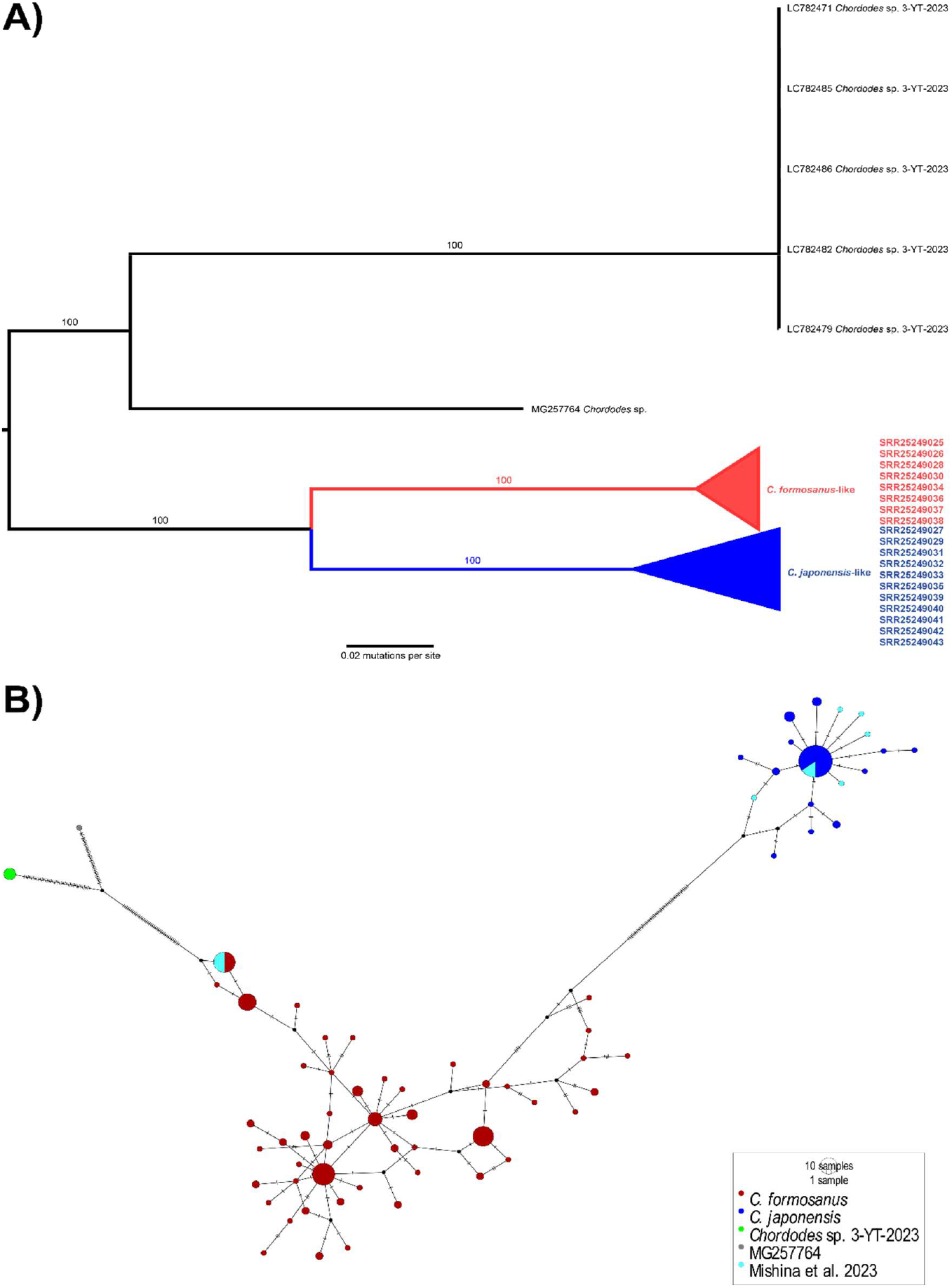
A) Collapsed phylogenetic tree based on COXI sequences, with also the SRA access number of the sequences from Mishina *et al*. (2023) and where they cluster. The unprocessed tree is available in Supplementary Data. B) Haplotype network based on the same COXI alignment as the tree.

### ddRAD + RNA-seq clustering analyses

After the filtering, 1,282 variant sites were retained from the 27 final individuals, with 12.92% missing data. In ipyrad analysis-tools, after the “minmap” step, around 1063 SNPs were retained, with 631 of them unlinked (Sup. Data). Using the unlinked SNPs, the PCA shows a split mainly in two groups; one of them is made up of the ddRAD Taiwanese samples and some from RNA-seq, mostly of them labelled as *C. formosanus* according to COXI, minus two individuals that were regarded as *C. japonensis*. The sequences in the other group were all regarded as *C. japonensis* according to COXI. There was also an individual that formed its own cluster (Fig. 3A). Such a pattern was also kept after the removal of the three individuals with exactly or more than 25% missing data, with one individual labelled as *C. japonensis* according to COXI still in the hybrid ddRAD/RNA-seq group that should represent *C. formosanus*, although the “rogue single-group” individual was not removed (Fig. 3B).

**Figure 3.**
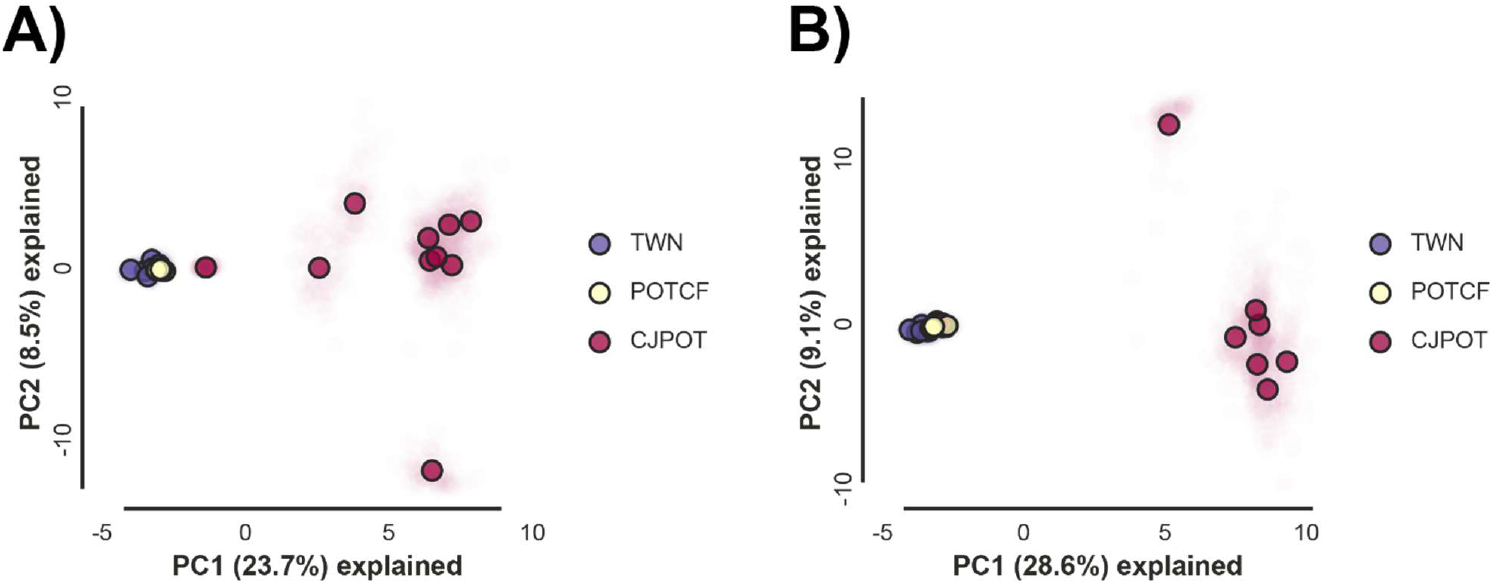
PCAs done with the ipyrad analysis-tools. A) PCA with the full 27 individuals’ dataset. B) PCA without individuals with exactly or more than 25% missing data. “TWN” = *C. formosanus* individuals from the Taiwanese ddRAD dataset. “POTCF” = specimens from Mishina *et al*. (2023) which were labelled as *C. formosanus* according to COXI. “CJPOT” = specimens from Mishina *et al*. (2023) which were labelled as *C. japonensis* according to COXI.

Using AIC and *snapclust* for both the full 27 individuals’ dataset and the one without the individuals with exactly or more than 25% missing data, it was shown how indeed the dataset was highly likely composed of two taxa (Sup. Fig. 1), although the cluster’s assignment differed compared to PCA with “find.clusters”: specifically, the samples SRR25249033 and SRR25249035 were found to be closer to the *C. formosanus*’s cluster (Sup. Fig. 2; Sup.

Data). SRR25249033 was found to be closer to the *C. formosanus*’s cluster also in the reduced dataset (Sup. Fig. 3; Data). K=2 was also the best cross-validation ADMIXTURE result (CV error=0.54822 and 0.56622 for the full dataset and the ones without individuals with exactly or more than 25% missing data, respectively: they were the lowest CV values for the 3 chosen K values per each dataset; Sup. Data). Said software’s results showed two clusters with both datasets that looked like the PCA groupings, although two individuals, SRR25249032 and SRR25249035, showed an 80 *C. japonensis*:20 *C. formosanus* ancestry ratio (Fig. 4). SRR25249031 showed an almost complete *C. japonensis* ancestry in the full dataset (Fig. 4A), while it showed roughly 95:5 ancestry ratio in the dataset with fewer individuals (Fig. 4B). The SRR25249029 and the SRR25249039 individuals, which were labelled as labelled as *C. japonensis* according to COXI, clustered 100% with *C. formosanus* specimens with both “find.clusters” and ADMIXTURE (Fig 4; Sup. Fig.s 2 and 3).

**Figure 4.**
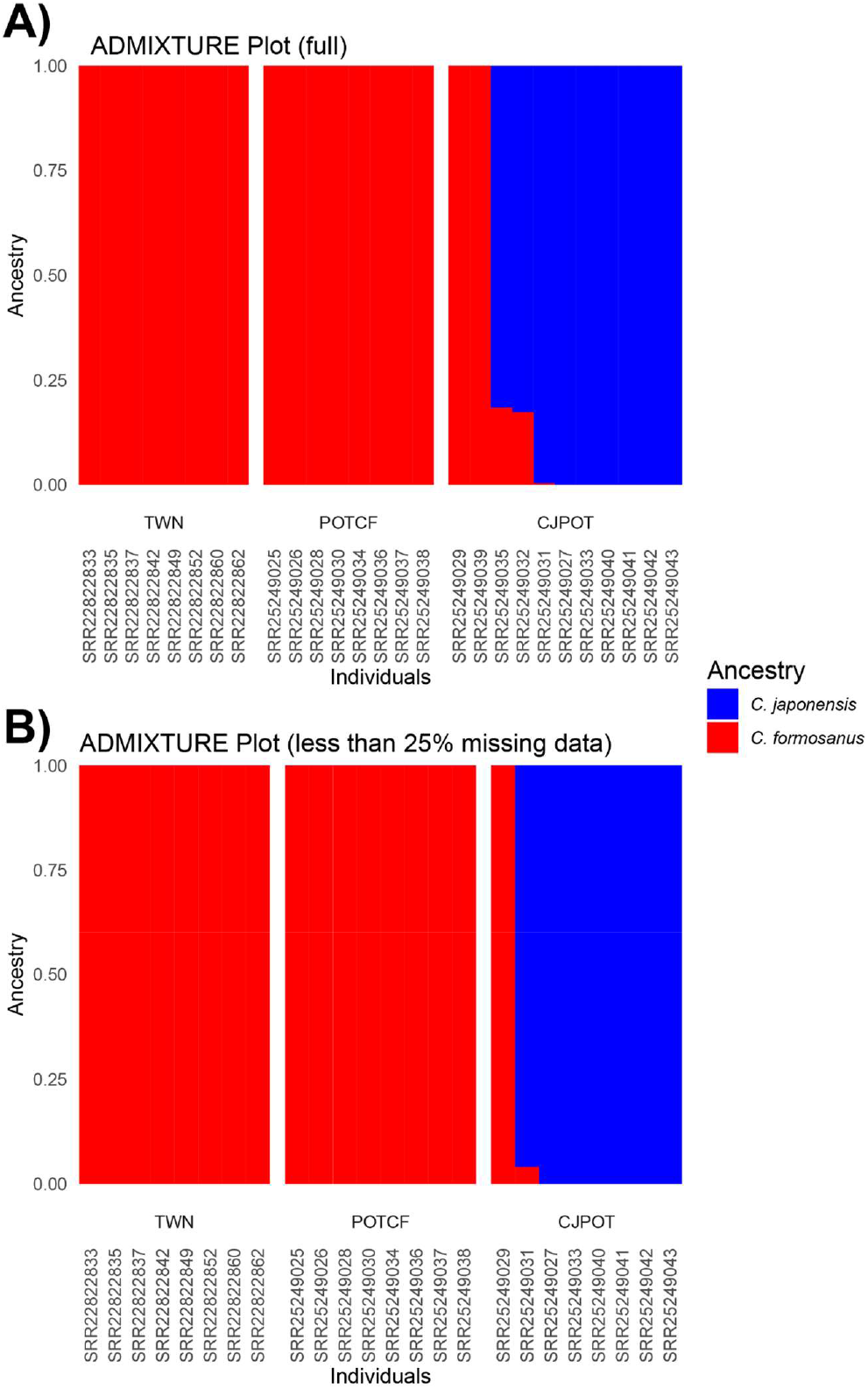
ADMIXTURE plots, with also individuals’ labeling. A) ADMIXTURE plots with the full 27 individuals’ dataset. B) ADMIXTURE plots without individuals with exactly or more than 25% missing data. “TWN” = *C. formosanus* individuals from the Taiwanese ddRAD dataset. “POTCF” = specimens from Mishina *et al*. (2023) which were labelled as *C. formosanus* according to COXI. “CJPOT” = specimens from Mishina *et al*. (2023) which were labelled as *C. japonensis* according to COXI.

### Demographic analyses

According to AIC, AIC weights and likelihood distribution, the *fastsimcoal* model that had different rates of gene flow between split and recent times was the most likely one, with both all the individuals and the SFS without the ones with exactly or more than 25% missing data (Table 2; Sup. Fig. 4 and 5; notice how the model with no geneflow has no data for the SFS with all the individuals, since one deme got extinct in all the 100 simulations with such model). Such events seem to be recent, given that the values of TDIV (first hybridization event) and CHANGM (time of second event) were around 358.81 (mean)±84.149 (SD) and 44.13±61.052, respectively, with the SFS with more individuals. Such values were around 759.42±140.571 for the former and around 94.5±40.554 for the latter with the SFS excluding the individuals with at least 25% missing data (Fig. 5; Sup. Data). The triangle plots show the presence of at least an individual (SRR25249031) who might be a backcross (Sup. Fig 6; Sup. Data).

**Table 2.**
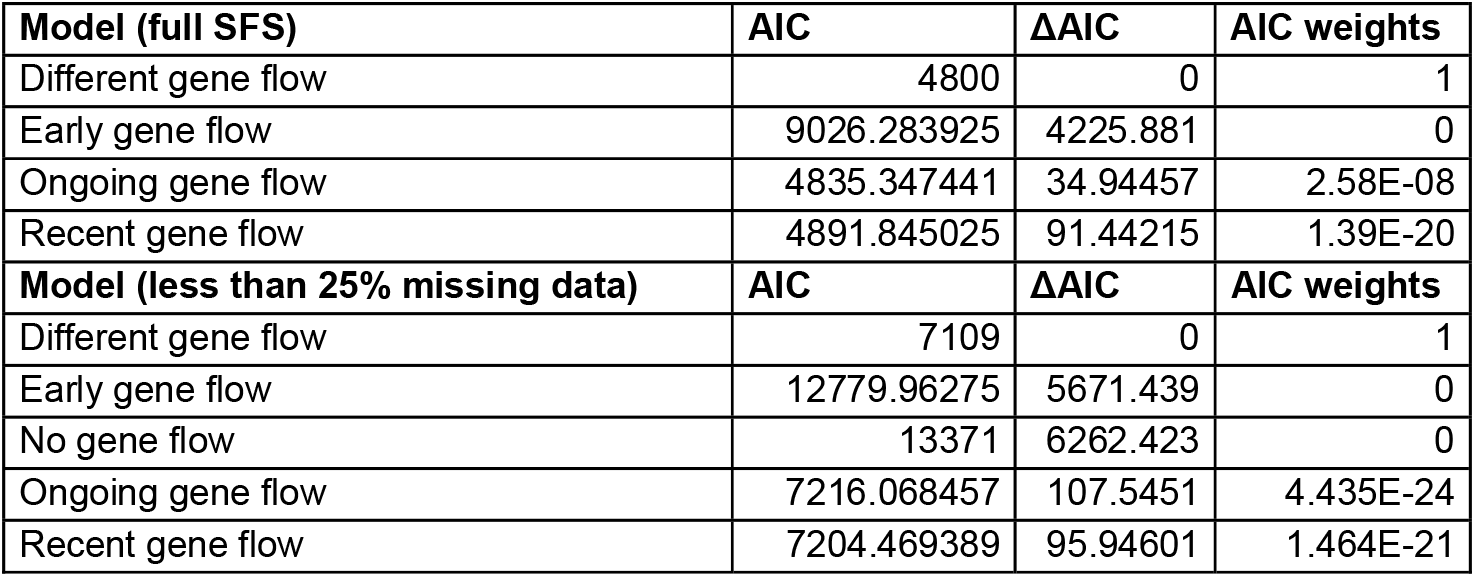
AICs of the *fastsimcoal* models. Notice that there is no data for the “no gene flow” model for the full SFS, since a deme got extinct in all the simulations.

| Model (full SFS) | AIC | ΔAIC | AIC weights |
| --- | --- | --- | --- |
| Different gene flow | 4800 | 0 | 1 |
| Early gene flow | 9026.283925 | 4225.881 | 0 |
| Ongoing gene flow | 4835.347441 | 34.94457 | 2.58E-08 |
| Recent gene flow | 4891.845025 | 91.44215 | 1.39E-20 |
| Model (less than 25% missing data) | AIC | ΔAIC | AIC weights |
| Different gene flow | 7109 | 0 | 1 |
| Early gene flow | 12779.96275 | 5671.439 | 0 |
| No gene flow | 13371 | 6262.423 | 0 |
| Ongoing gene flow | 7216.068457 | 107.5451 | 4.435E-24 |
| Recent gene flow | 7204.469389 | 95.94601 | 1.464E-21 |

**Figure 5.**
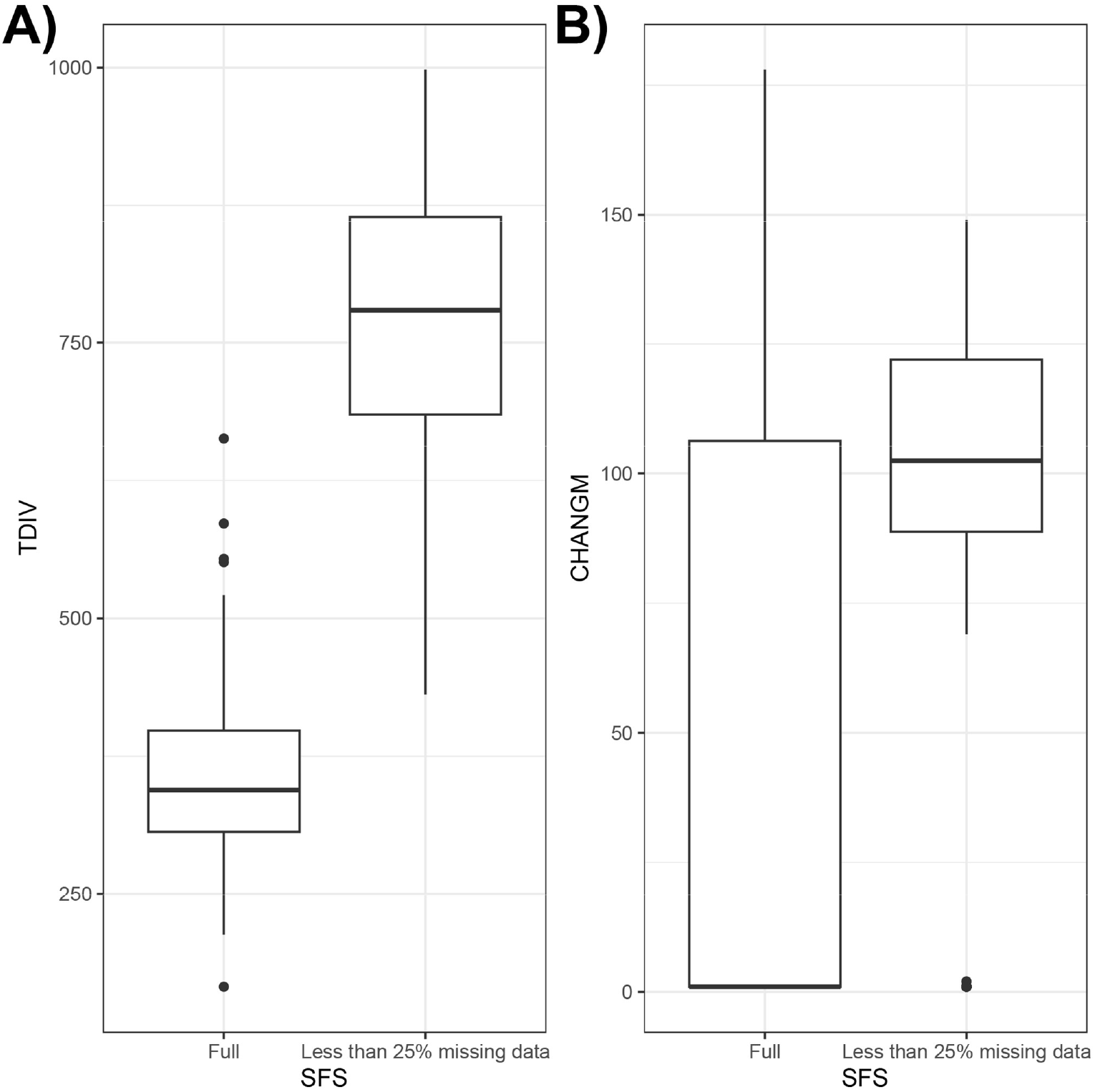
A) Boxplots of the values of TDIV (first hybridization event). B) Boxplots of the values of CHANGM (second hybridization event). All the boxplots were based on full or partial (without the individuals with at least 25% missing data) SFS.

## Discussion

In this paper, I report a species misidentification from a previous paper, since what was reported as *Chordodes fukuii* in Mishina *et al*. (2023) seems to consist of two actual groups. One group corresponds to *C. formosanus*, which is known from Japan and Taiwan (Chiu, 2017 and references therein). The other group seems to correspond to *C. japonensis*, which is known from Japan and Korea (Schmidt-Rhaesa, 2004 and references therein), given the high COXI similarity and the clustering with specimens labelled as such using the barcoding gene.

Interestingly, all the *Chordodes* specimens from Mishina *et al*. (2023) come from a single host species, the narrow-winged mantis (*Tenodera angustipennis*). Chiu (2017) reports how no *C. formosanus* specimen was ever reported from *Tenodera* species. However, reports of *C. formosanus* from *T. angustipennis* in Japan increased recently (Kuroda *et al*., 2024; Tani *et al*., 2024), showing how this hairworm is more flexible in host choice than what it was though. Lack of reports from Taiwanese *T. angustipennis* might be caused by undersampling or preference for the giant Taiwanese mantis (*Titanodula formosana*, formerly regarded as *Hierodula formosana*; see De Vivo *et al*. 2023a and references therein), which is the most used host in Taiwan (Chiu, 2017 and references therein). Albeit *C. formosanus* is usually found way more often in giant Asian mantises (subfamily Hierodulinae: Chiu et al., 2011; Tani *et al*., 2024) and *C. japonensis* is usually associated with *Tenodera* species (Inoue, 1952; Chiu *et al*., 2011; Tani *et al*., 2024), there are reports of both species in different mantises and even katydids (family Tettigoniidae, albeit those might be rarely used and generally not preferred; Chiu, 2017; Chiu *et al*., 2017; Tani *et al*., 2024). Furthermore, *C. fukuii* also uses *Tenodera* species as definitive hosts (Inoue, 1952). Given the following considerations, species identification in horsehair worms cannot be based only by checking from which definitive host they emerge, albeit host choice might be a phylogenetic signal at genus level.

Generally, as said in the Introduction, species identification in horsehair worms is usually regarded as difficult due to the paucity of morphological characters useful for distinguishing clades, which often are visible only through SEM and/or DIC microscopy (Schmidt-Rhaesa, 2013). Chiu *et al*. (2011) highlighted similarities and dissimilarities between *C. formosanus* and *C. japonensis*, noticing how the former has significantly longer crowned areoles’ filaments in female specimens and lacks long-crowned areoles in male individuals. Schmidt-Rhaesa (2004) reports the lack of “characteristic clusters” of areoles in *C. fukuii* compared to *C. japonensis*. Given that dissimilarities are minimal, some previous work used COXI without SEM for species ID in East Asian *Chordodes*, particularly with damaged specimens (De Vivo *et al*., 2023a; Kuroda *et al*., 2024; Tani *et al*., 2024). In Chiu *et al*. (2017), some *Chordodes* specimens were recognized as *C. formosanus* by barcoding after noticing morphological abnormalities, potentially caused by development in the “wrong” definitive host (katydids), which made morphological species identification challenging. De Vivo *et al*. (2023a) showed how COXI works well for species ID in *C. formosanus*, given the matching ID from such barcoding gene with the one from previous studies (Chiu *et al*., 2011 & 2017) and the lack of population structure in Taiwanese specimens inferred by ddRAD.

It is highly likely the specimens from Mishina *et al*. (2023) were not correctly processed for morphological taxonomic ID due to the need of high-quality RNA from extraction, since the RNA was extracted from the worms’ whole body. Additionally, no *C. fukuii* COXI data is currently available, making ID by barcoding almost impossible for such species; for example, Kuroda *et al*. (2024) barcoded some *Chordodes* specimens and were not able to assign some of these individuals to a species due to lack of data in public repositories and no usage of microscopy for taxonomic identification, albeit the specimens might belong to either *C. fukuii* or *C. silvestri* (see Inoue 1952 and Schmidt-Rhaesa 2004 for the presence of *Chordodes* species in Japan). That being said, the results from Mishina *et al*. (2023) become stronger with this paper, even if the species were misidentified; specifically, worms clustered together according to the manipulation stage in the previous publication, even if the species ID is different among worms, showing potential similarities in host-parasite interactions.

Even with molecular data, some individuals clearly labelled as *C. japonensis* with COXI were not assigned with certainty to clusters in the ddRAD-RNAseq dataset. The presence and the location of missing data might be the reason why clustering was unclear, since it is known that those factors can cause issues in identifying groups with PCA and similar methods (Yi & Latch, 2022). That being said, one specimen (SRR25249029) showed 100% mito-nuclear discordance, i.e. it 100% belonged to *C. japonensis* with COXI and *C. formosanus* with more data in all the possible datasets (complete and the ones without individuals with exactly more than 25% missing data). SRR25249039 showed a similar pattern (albeit such specimen had at least 25% missing data, making the result more questionable), and three other specimens (SRR25249031, SRR25249032 and SRR25249035) showed a mixed ancestry in the ddRAD-RNA datasets, albeit mostly derived from *C. japonensis*. This might be caused by missing data in certain locations (particularly for SRR25249032 and SRR25249035, since they had exactly 25% missing data), but it could also be a sign of previous introgression by hybridization, which is reported in other helminthic taxa (see Detwiler & Criscione 2010). A potential hybridization scenario, with probably two hybridization events with different gene flow between the species, was also regarded as the most likely model by *fastsimcoal* in this study and triangle plots show a potential backcross individual, SRR25249031.

Such introgression, if confirmed, seems to be biased, given the presence of a mitochondrial lineage from only one species (*C. japonensis*) in these potentially mixed individuals. Therefore, potential recent or ancient hybridization might come from male *C. formosanus* breeding with female *C. japonensis*. Females mating with heterospecific males is a known phenomenon (see “Box 1” in Pfennig 2021), albeit hybridization in general is not reported in hairworms yet. If such phenomenon is confirmed, it might be caused by the skewed sex ratio in hairworms (Schmidt-Rhaesa, 2013 and references therein) or lower levels of survival in hybrids from male *C. japonensis* and female *C. formosanus*, following the Darwin’s Corollary to Haldane’s Rule (Turelli & Moyle, 2007). However, another potential valid explanation for this pattern might be incomplete lineage sorting (ILS), caused by ancestral genetic variations that persisted through speciation or, as said earlier, missing data in certain locations. ILS can lead to conflicting topologies, and it is often hard to distinguish it from recent or past introgression (Detwiler & Criscione, 2010).

Additionally, it must be noted that demographic simulations may have biases when events are recent (Shen *et al*., 2025) or when they do not account for multiple demographic events (e.g., Momigliano *et al*. 2021). Therefore, the recent estimated ages for hybridization according to bootstrap replicates should not be taken as absolute truth. It might be possible that such events are more ancient than what the simulations suggest, although here it may be tough to quantify how much. Specifically, it is possible that both species are more widespread in Asia than is thought: currently, *C. japonensis* is also reported from Korea (Baek, 1993), while *C. formosanus* is widespread in Taiwan, including the volcanic island of Lyudao (Chiu, 2017; De Vivo *et al*., 2023a). If their geographic origin is not clear, it is challenging to make a potential link between ancient events (e.g., geological ones like the uplift of the island of Taiwan or the formation of the Japanese archipelago) and their population demography. That being said, all the top *fastsimcoal* models used here show that hybridization indeed happened, albeit such results may be influenced by some biases highlighted in the previous sections.

All the potential explanations for the mito-nuclear discordance shown here might be proved with data from an outgroup congeneric species and further analyses like ABBA-BABA tests. At the time of the last GenBank check (June 24, 2025), the closest species to *Chordodes* with SRA data on GenBank was highly likely *Parachordes pustulosus* from Europe (Nikolaeva *et al*., 2023), which however is considered closer based on single gene data and morphology (Schmidt-Rhaesa, 2013; Nikolaeva *et al*., 2023). Furthermore, it may be possible that *Parachordodes* could be phylogenetically far from *Chordodes*. Specifically, although morphology is extremely conserved among clades, Nematomorpha is an ancient group, which probably emerged between mid-Cambrian and mid-Cretaceous (Howard *et al*., 2022).

Freshwater hairworms definitively were already present 110/90 millions of years ago (Poinar & Buckley, 2006; Howard *et al*., 2022); given this, choosing a species from another genera as an outgroup might be risky, given the ancient age of the group, the unresolved phylogenetic questions regarding relationships at different clade levels (family and genus) and the potential ancient split among genera.

Given this, I argue that several questions and doubts could be answered or clarified by sequencing a genome of an outgroup *Chordodes* species and using it as a reference for assembly. Specifically, it has been shown how reference genomes allow to distinguish hybridization from ILS and check for linkage blocks, which can lead to estimation of linkage disequilibrium and better data filtering, while also visualizing potential recombination areas (Theissinger *et al*., 2023 and references therein). Furthermore, more loci could be recovered through a reference genome and it can help to distinguish recent demographic events from past ones better (Shen *et al*., 2025); particularly the last point is critical, since hybridization events seem to be recent according to the *fastsimcoal* simulations in this study. Therefore, a genome assembly of a *Chordodes* species could be important for understanding hairworm ecology better and their genomics.

## Conclusions

I generated COXI data from previously released RNAseq data and also analyzed a species misidentification in horsehair worms, highlighting potential introgression between *Chordodes* species. Furthermore, I also show each needed step for doing this kind of analyses, which can be replicated and used for studies regarding any other species with data available. This paper also shows the need of a reference genome for *Chordodes* species for understanding if there is any current or ancient hybridization in Nematomorpha, one of the most understudied animal groups, and the need in increasing the number of available barcodes for morphologically recognized species.

## Supporting information

Supplementary Materials (Tables and Images) for the study

## Acknowledgments

I would like to thank Nico Fuhrmann (Trier University, Trier, Germany), whose help and discussion regarding the assembling of the ddRAD-RNAseq dataset was pivotal for the study. I would also like to thank Vladislav “Uładź” Ivanov (University of Rostock, Rostock, Germany) for discussion regarding the ddRAD-RNAseq dataset and Jen-Pan Huang (Academia Sinica, Taipei, Taiwan) for suggestions on how to evaluate potential hybridization.

## Data Availability

Data and scripts are available on Zenodo (doi: 10.5281/zenodo.18415952) at https://doi.org/10.5281/zenodo.18415952.

