## Supplementary Materials (Tables and Images) for the study for "How many are you? Open data and bioinformatics reveal species misidentification and potential introgression in *Chordodes* (Phylum Nematomorpha)"

### Supplementary Tables and figures

### Supplementary Tables

| RNA SRA ID | Top match on Genbank | Species ID | % ID (BLASTn) |
|---|---|---|---|
| SRR25249025 | LC782483 | *C. formosanus* | 100 |
| SRR25249026 | LC782483 | *C. formosanus* | 100 |
| SRR25249027 | LC776436 | *C. japonensis* | 99.696 |
| SRR25249028 | LC782483 | *C. formosanus* | 100 |
| SRR25249029 | LC776416 | *C. japonensis* | 100 |
| SRR25249030 | LC782483 | *C. formosanus* | 100 |
| SRR25249031 | LC776441 | *C. japonensis* | 100 |
| SRR25249032 | LC782455 | *C. japonensis* | 100 |
| SRR25249033 | LC776441 | *C. japonensis* | 100 |
| SRR25249034 | LC782483 | *C. formosanus* | 100 |
| SRR25249035 | LC776436 | *C. japonensis* | 100 |
| SRR25249036 | LC782483 | *C. formosanus* | 100 |
| SRR25249037 | LC782483 | *C. formosanus* | 100 |
| SRR25249038 | LC782483 | *C. formosanus* | 100 |
| SRR25249039 | LC776441 | *C. japonensis* | 100 |
| SRR25249040 | LC776416 | *C. japonensis* | 99.848 |
| SRR25249041 | LC776418 | *C. japonensis* | 99.68 |
| SRR25249042 | LC776436 | *C. japonensis* | 99.696 |
| SRR25249043 | LC776436 | *C. japonensis* | 99.848 |

**Sup. Table 1.** BLASTn results for the COXI extracted from RNA SRAs of Mishina *et al*. (2023).

| Accession number | Source |
|---|---|
| HM044104 | Chiu *et al*. (2011) |
| HM044105 | Chiu *et al*. (2011) |
| HM044106 | Chiu *et al*. (2011) |
| HM044107 | Chiu *et al*. (2011) |
| HM044108 | Chiu *et al*. (2011) |
| HM044109 | Chiu *et al*. (2011) |
| HM044110 | Chiu *et al*. (2011) |
| HM044111 | Chiu *et al*. (2011) |
| HM044112 | Chiu *et al*. (2011) |
| HM044113 | Chiu *et al*. (2011) |
| HM044114 | Chiu *et al*. (2011) |
| HM044115 | Chiu *et al*. (2011) |
| HM044116 | Chiu *et al*. (2011) |
| HM044117 | Chiu *et al*. (2011) |
| HM044118 | Chiu *et al*. (2011) |
| HM044119 | Chiu *et al*. (2011) |
| HM044120 | Chiu *et al*. (2011) |
| HM044121 | Chiu *et al*. (2011) |
| HM044122 | Chiu *et al*. (2011) |
| HM044123 | Chiu *et al*. (2011) |
| HM044124 | Chiu *et al*. (2011) |
| HM044125 | Chiu *et al*. (2011) |
| HM044126 | Chiu *et al*. (2011) |
| HM044127 | Chiu *et al*. (2011) |
| HM044128 | Chiu *et al*. (2011) |
| HQ322115 | Chiu *et al*. (2011) |
| HQ322116 | Chiu *et al*. (2011) |
| JF808194 | Chiu *et al*. (2011) |
| JF808195 | Chiu *et al*. (2011) |
| JF808196 | Chiu *et al*. (2011) |
| JF808197 | Chiu *et al*. (2011) |
| JF808198 | Chiu *et al*. (2011) |
| JF808199 | Chiu *et al*. (2011) |
| JF808200 | Chiu *et al*. (2011) |
| JF808201 | Chiu *et al*. (2011) |
| JF808202 | Chiu *et al*. (2011) |
| JF808203 | Chiu *et al*. (2011) |
| JF808204 | Chiu *et al*. (2011) |
| JF808205 | Chiu *et al*. (2011) |
| JF808206 | Chiu *et al*. (2011) |
| KU509484 | Chiu *et al*. (2015) |
| KU509485 | Chiu *et al*. (2015) |

| | |
|---|---|
| KU509486 | Chiu *et al*. (2015) |
| KU509487 | Chiu *et al*. (2015) |
| KU509488 | Chiu *et al*. (2015) |
| KX591949 | Chiu *et al*. (2015) |
| KX591950 | Chiu *et al*. (2015) |
| KX591951 | Chiu *et al*. (2015) |
| KX591952 | Chiu *et al*. (2015) |
| KX591953 | Chiu *et al*. (2015) |
| LC776384 | Tani *et al*. (2024) |
| LC776385 | Tani *et al*. (2024) |
| LC776386 | Tani *et al*. (2024) |
| LC776387 | Tani *et al*. (2024) |
| LC776388 | Tani *et al*. (2024) |
| LC776389 | Tani *et al*. (2024) |
| LC776390 | Tani *et al*. (2024) |
| LC776391 | Tani *et al*. (2024) |
| LC776392 | Tani *et al*. (2024) |
| LC776393 | Tani *et al*. (2024) |
| LC776394 | Tani *et al*. (2024) |
| LC776395 | Tani *et al*. (2024) |
| LC776396 | Tani *et al*. (2024) |
| LC776397 | Tani *et al*. (2024) |
| LC776398 | Tani *et al*. (2024) |
| LC776399 | Tani *et al*. (2024) |
| LC776400 | Tani *et al*. (2024) |
| LC776401 | Tani *et al*. (2024) |
| LC776402 | Tani *et al*. (2024) |
| LC776403 | Tani *et al*. (2024) |
| LC776404 | Tani *et al*. (2024) |
| LC776405 | Tani *et al*. (2024) |
| LC776406 | Tani *et al*. (2024) |
| LC776407 | Tani *et al*. (2024) |
| LC776408 | Tani *et al*. (2024) |
| LC776409 | Tani *et al*. (2024) |
| LC776410 | Tani *et al*. (2024) |
| LC776411 | Tani *et al*. (2024) |
| LC776412 | Tani *et al*. (2024) |
| LC776413 | Tani *et al*. (2024) |
| LC776414 | Tani *et al*. (2024) |
| LC776415 | Tani *et al*. (2024) |
| LC776416 | Tani *et al*. (2024) |
| LC776417 | Tani *et al*. (2024) |
| LC776418 | Tani *et al*. (2024) |
| LC776419 | Tani *et al*. (2024) |

| | |
|---|---|
| LC776420 | Tani *et al*. (2024) |
| LC776421 | Tani *et al*. (2024) |
| LC776422 | Tani *et al*. (2024) |
| LC776423 | Tani *et al*. (2024) |
| LC776424 | Tani *et al*. (2024) |
| LC776425 | Tani *et al*. (2024) |
| LC776426 | Tani *et al*. (2024) |
| LC776427 | Tani *et al*. (2024) |
| LC776428 | Tani *et al*. (2024) |
| LC776429 | Tani *et al*. (2024) |
| LC776430 | Tani *et al*. (2024) |
| LC776431 | Tani *et al*. (2024) |
| LC776432 | Tani *et al*. (2024) |
| LC776433 | Tani *et al*. (2024) |
| LC776434 | Tani *et al*. (2024) |
| LC776435 | Tani *et al*. (2024) |
| LC776436 | Tani *et al*. (2024) |
| LC776437 | Tani *et al*. (2024) |
| LC776438 | Tani *et al*. (2024) |
| LC776439 | Tani *et al*. (2024) |
| LC776440 | Tani *et al*. (2024) |
| LC776441 | Tani *et al*. (2024) |
| LC776442 | Tani *et al*. (2024) |
| LC776443 | Tani *et al*. (2024) |
| LC776444 | Tani *et al*. (2024) |
| LC776445 | Tani *et al*. (2024) |
| LC776446 | Tani *et al*. (2024) |
| LC776447 | Tani *et al*. (2024) |
| LC776448 | Tani *et al*. (2024) |
| LC776449 | Tani *et al*. (2024) |
| LC776450 | Tani *et al*. (2024) |
| LC776451 | Tani *et al*. (2024) |
| LC776452 | Tani *et al*. (2024) |
| LC776453 | Tani *et al*. (2024) |
| LC776454 | Tani *et al*. (2024) |
| LC776455 | Tani *et al*. (2024) |
| LC776456 | Tani *et al*. (2024) |
| LC776457 | Tani *et al*. (2024) |
| LC776458 | Tani *et al*. (2024) |
| LC782468 | Kuroda *et al*. (2024) |
| LC782469 | Kuroda *et al*. (2024) |
| LC782470 | Kuroda *et al*. (2024) |
| LC782471 | Kuroda *et al*. (2024) |
| LC782472 | Kuroda *et al*. (2024) |

| | |
|---|---|
| LC782473 | Kuroda *et al*. (2024) |
| LC782474 | Kuroda *et al*. (2024) |
| LC782475 | Kuroda *et al*. (2024) |
| LC782476 | Kuroda *et al*. (2024) |
| LC782477 | Kuroda *et al*. (2024) |
| LC782478 | Kuroda *et al*. (2024) |
| LC782479 | Kuroda *et al*. (2024) |
| LC782480 | Kuroda *et al*. (2024) |
| LC782481 | Kuroda *et al*. (2024) |
| LC782482 | Kuroda *et al*. (2024) |
| LC782483 | Kuroda *et al*. (2024) |
| LC782484 | Kuroda *et al*. (2024) |
| LC782485 | Kuroda *et al*. (2024) |
| LC782486 | Kuroda *et al*. (2024) |
| MG257764 | Mikhailov *et al*. (2019) |
| OQ121048 | De Vivo *et al.* (2023a) |
| OQ121049 | De Vivo *et al.* (2023a) |
| OQ121050 | De Vivo *et al*. (2023a) |
| OQ121051 | De Vivo *et al*. (2023a) |
| OQ121052 | De Vivo *et al*. (2023a) |
| OQ121053 | De Vivo *et al*. (2023a) |
| OQ121054 | De Vivo *et al*. (2023a) |
| OQ121055 | De Vivo *et al*. (2023a) |
| OQ121056 | De Vivo *et al*. (2023a) |
| OQ121057 | De Vivo *et al*. (2023a) |
| OQ121058 | De Vivo *et al*. (2023a) |
| OQ121059 | De Vivo *et al*. (2023a) |
| OQ121060 | De Vivo *et al*. (2023a) |
| OQ121061 | De Vivo *et al*. (2023a) |
| OQ121062 | De Vivo *et al*. (2023a) |
| OQ121063 | De Vivo *et al*. (2023a) |
| OQ121064 | De Vivo *et al*. (2023a) |
| OQ121065 | De Vivo *et al*. (2023a) |
| OQ121066 | De Vivo *et al*. (2023a) |
| OQ121067 | De Vivo *et al*. (2023a) |
| OQ121068 | De Vivo *et al*. (2023a) |
| OQ121069 | De Vivo *et al*. (2023a) |
| OQ121070 | De Vivo *et al*. (2023a) |
| OQ121071 | De Vivo *et al*. (2023a) |
| OQ121072 | De Vivo *et al*. (2023a) |
| OQ121073 | De Vivo *et al*. (2023a) |
| OQ121074 | De Vivo *et al*. (2023a) |
| OQ121075 | De Vivo *et al*. (2023a) |
| OQ121076 | De Vivo *et al*. (2023a) |

| | |
|---|---|
| OQ121077 | De Vivo *et al*. (2023a) |
| OQ121078 | De Vivo *et al*. (2023a) |
| OQ121079 | De Vivo *et al*. (2023a) |
| OQ121080 | De Vivo *et al*. (2023a) |
| OQ121081 | De Vivo *et al*. (2023a) |
| OQ121082 | De Vivo *et al*. (2023a) |
| OQ121083 | De Vivo *et al*. (2023a) |

**Sup. Table 2.** GenBank accession numbers of the downloaded COXI sequences.

| SRA |
|---|
| SRR22822824 |
| SRR22822825 |
| SRR22822827 |
| SRR22822833 |
| SRR22822834 |
| SRR22822835 |
| SRR22822836 |
| SRR22822837 |
| SRR22822838 |
| SRR22822839 |
| SRR22822842 |
| SRR22822843 |
| SRR22822846 |
| SRR22822847 |
| SRR22822848 |
| SRR22822849 |
| SRR22822850 |
| SRR22822851 |
| SRR22822852 |
| SRR22822854 |
| SRR22822858 |
| SRR22822859 |
| SRR22822860 |
| SRR22822862 |
| SRR22822864 |
| SRR22822865 |
| SRR22822866 |

**Sup. Table 3.** SRAs of all the downloaded ddRADseq data.

| Sample | Missing data |
|---|---|
| SRR25249041 | 0.01 |
| SRR25249042 | 0.01 |
| SRR25249040 | 0.01 |
| SRR25249027 | 0.02 |
| SRR25249038 | 0.02 |
| SRR25249043 | 0.02 |
| SRR25249037 | 0.03 |
| SRR25249028 | 0.04 |
| SRR25249029 | 0.05 |
| SRR25249031 | 0.06 |
| SRR22822835 | 0.07 |
| SRR22822860 | 0.08 |
| SRR22822862 | 0.1 |
| SRR22822833 | 0.1 |
| SRR25249025 | 0.12 |
| SRR25249026 | 0.12 |
| SRR25249036 | 0.13 |
| SRR22822852 | 0.15 |
| SRR22822842 | 0.17 |
| SRR25249030 | 0.17 |
| SRR25249034 | 0.18 |
| SRR25249033 | 0.19 |
| SRR22822849 | 0.2 |
| SRR22822837 | 0.21 |
| SRR25249032 | 0.25 |
| SRR25249035 | 0.25 |
| SRR25249039 | 0.27 |

**Sup. Table 4.** The amount of missing data in the final VCF assembly according to ipyrad analysis-tools.

A)

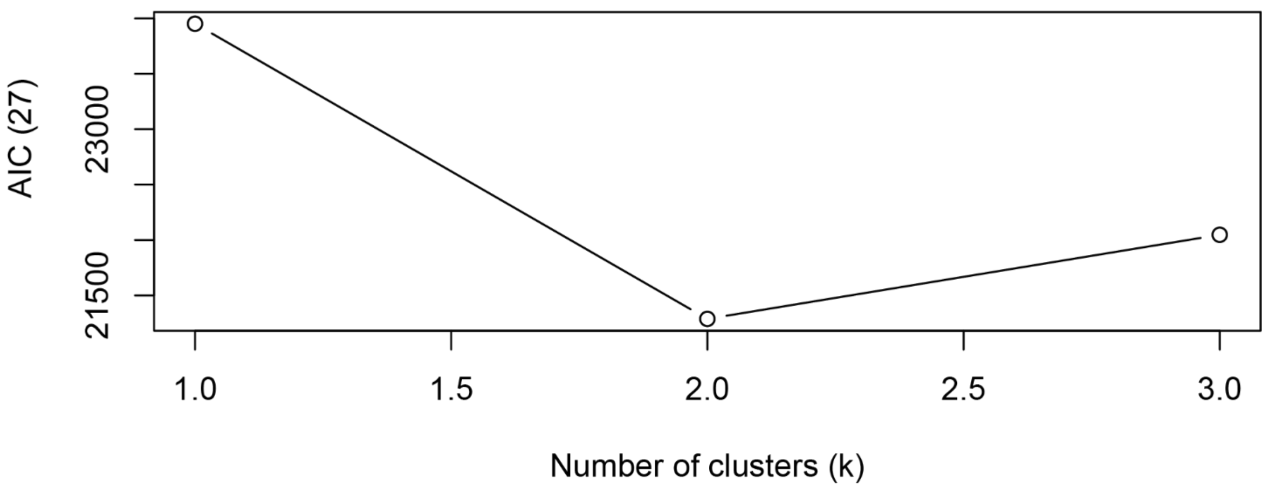


B)

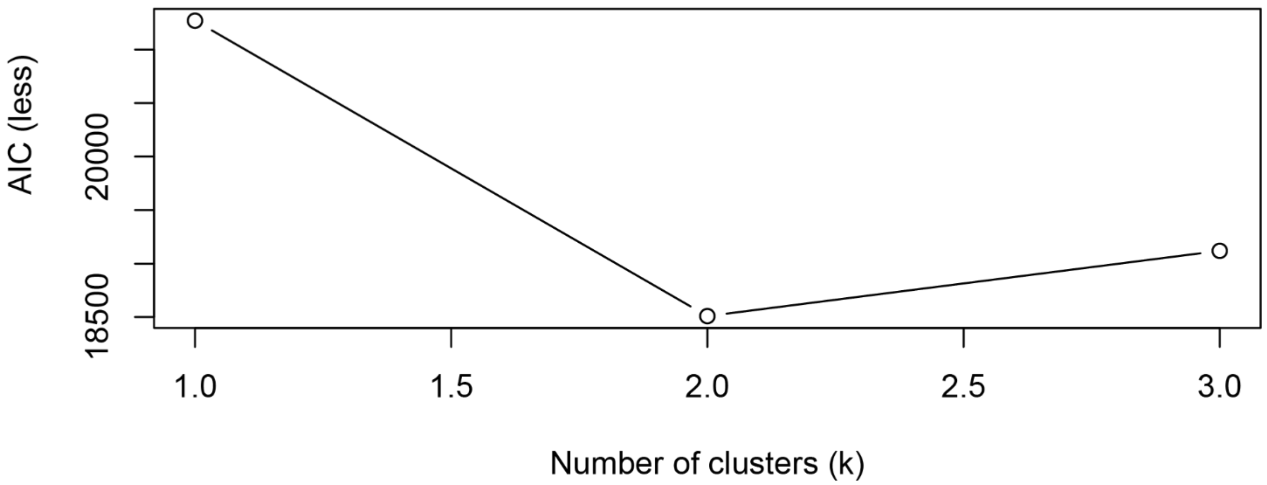


**Sup. Fig 1.** AIC calculated by snapclust. A) AIC for the whole dataset. B) AIC for the individuals with less than 25% missing data.

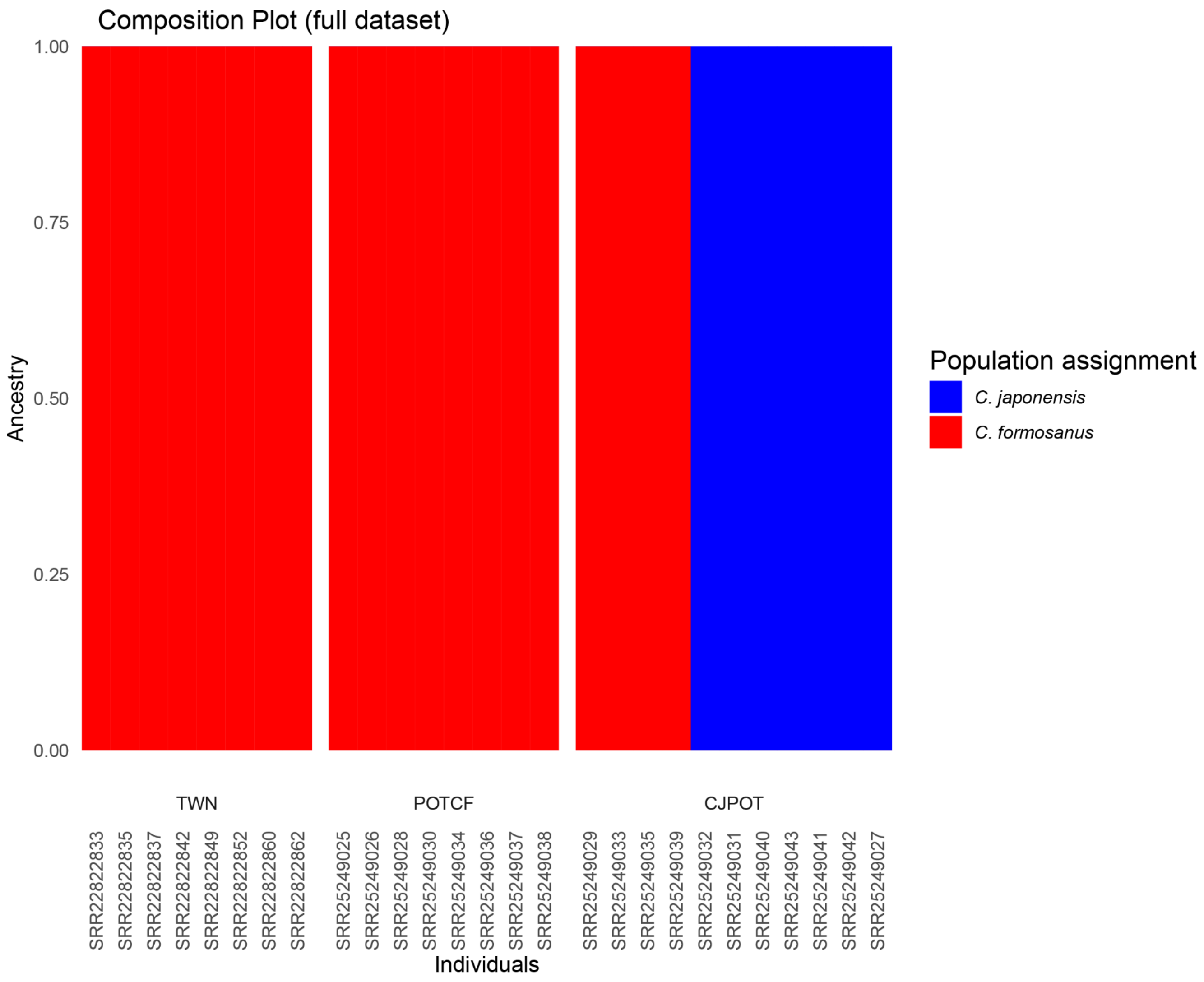


**Sup. Fig 2.** Composition plot from “find.clusters”, with the full dataset.

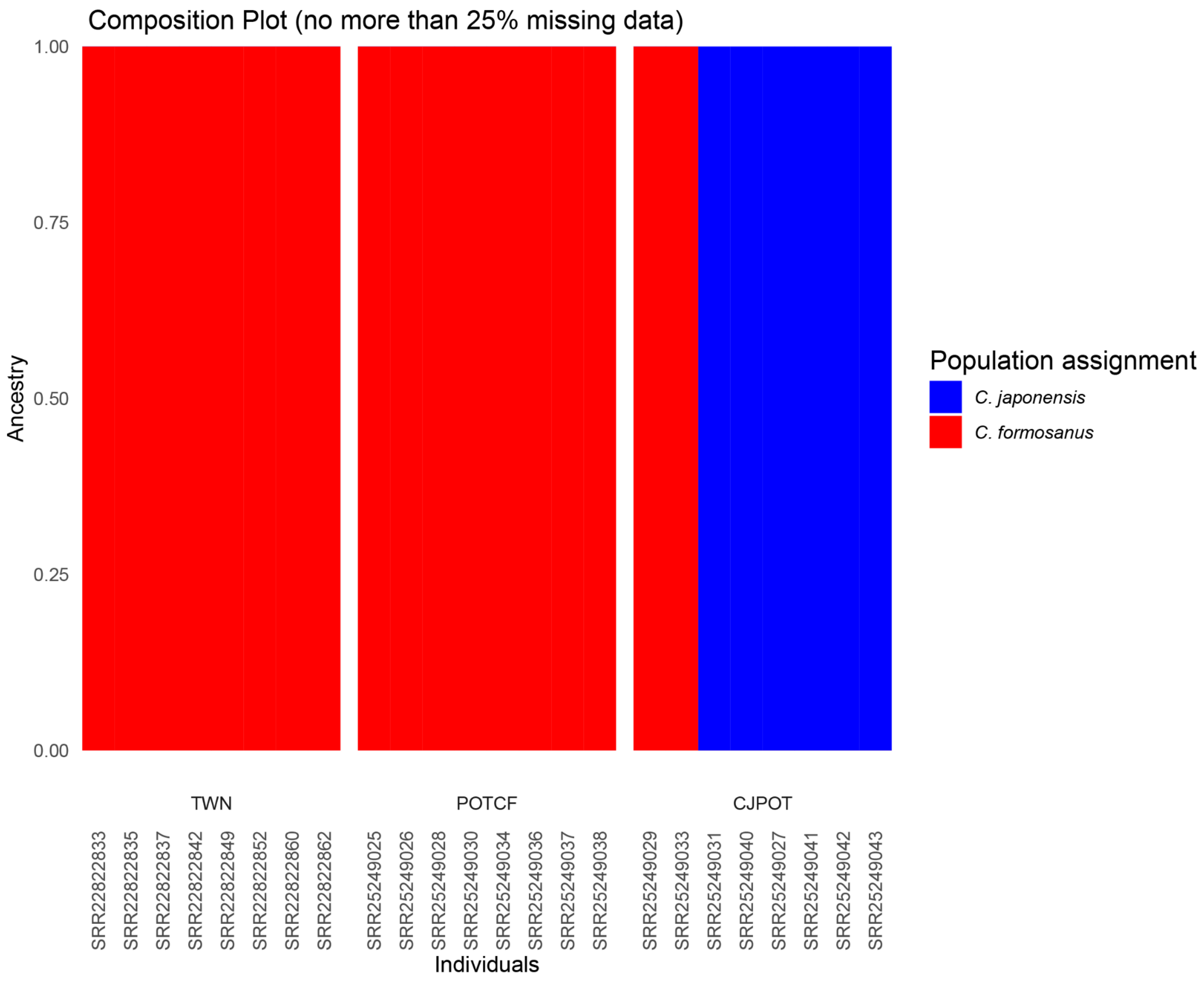


**Sup. Fig. 3.** Composition plot from “find.clusters”, with individuals with less than 25% missing data.

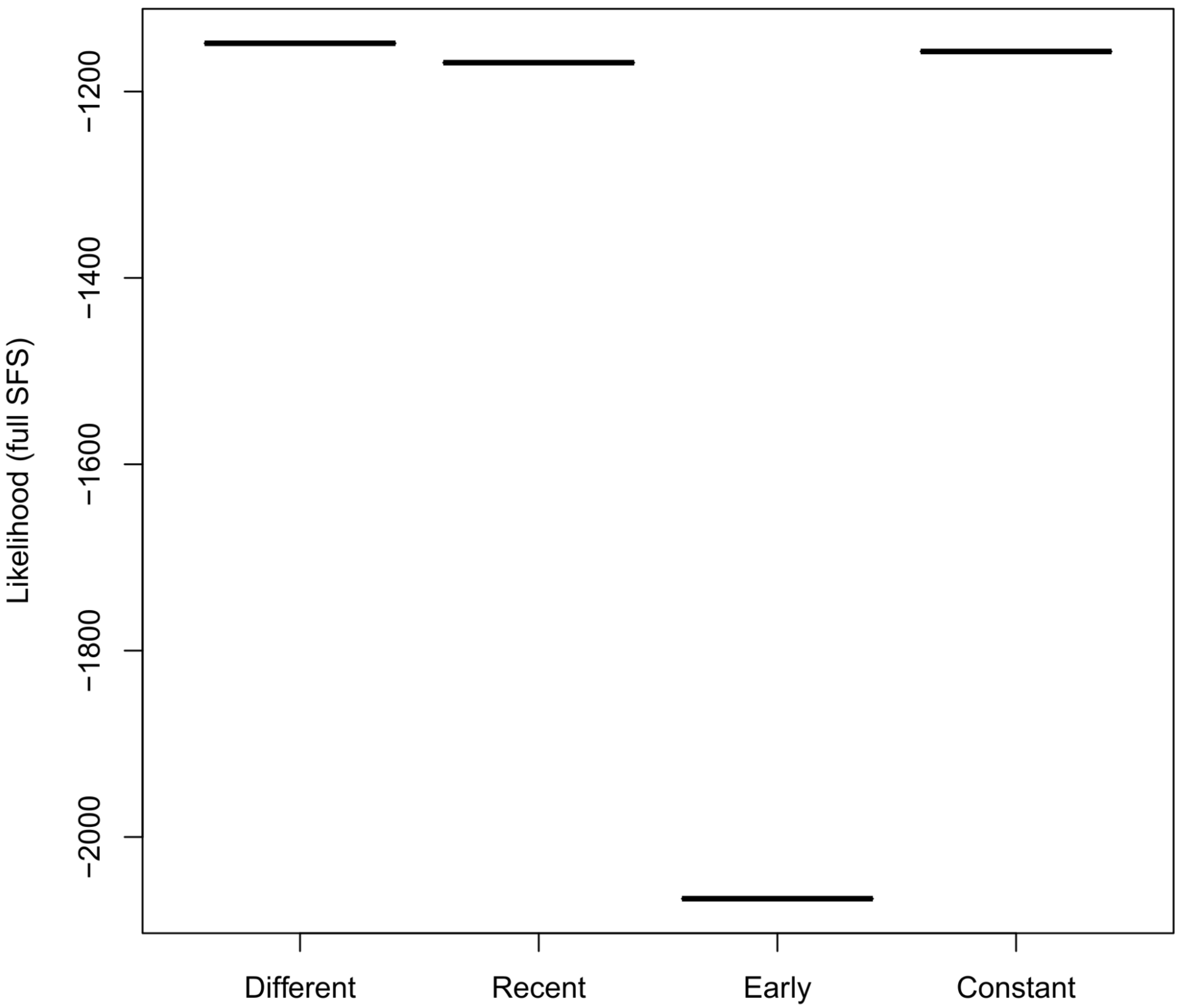


**Sup. Fig. 4.** Likelihood distribution for the *fastsimcoal* models with the full SFS. Notice that there is no data for the “no gene flow” model, since a deme got extinct in all the simulations.

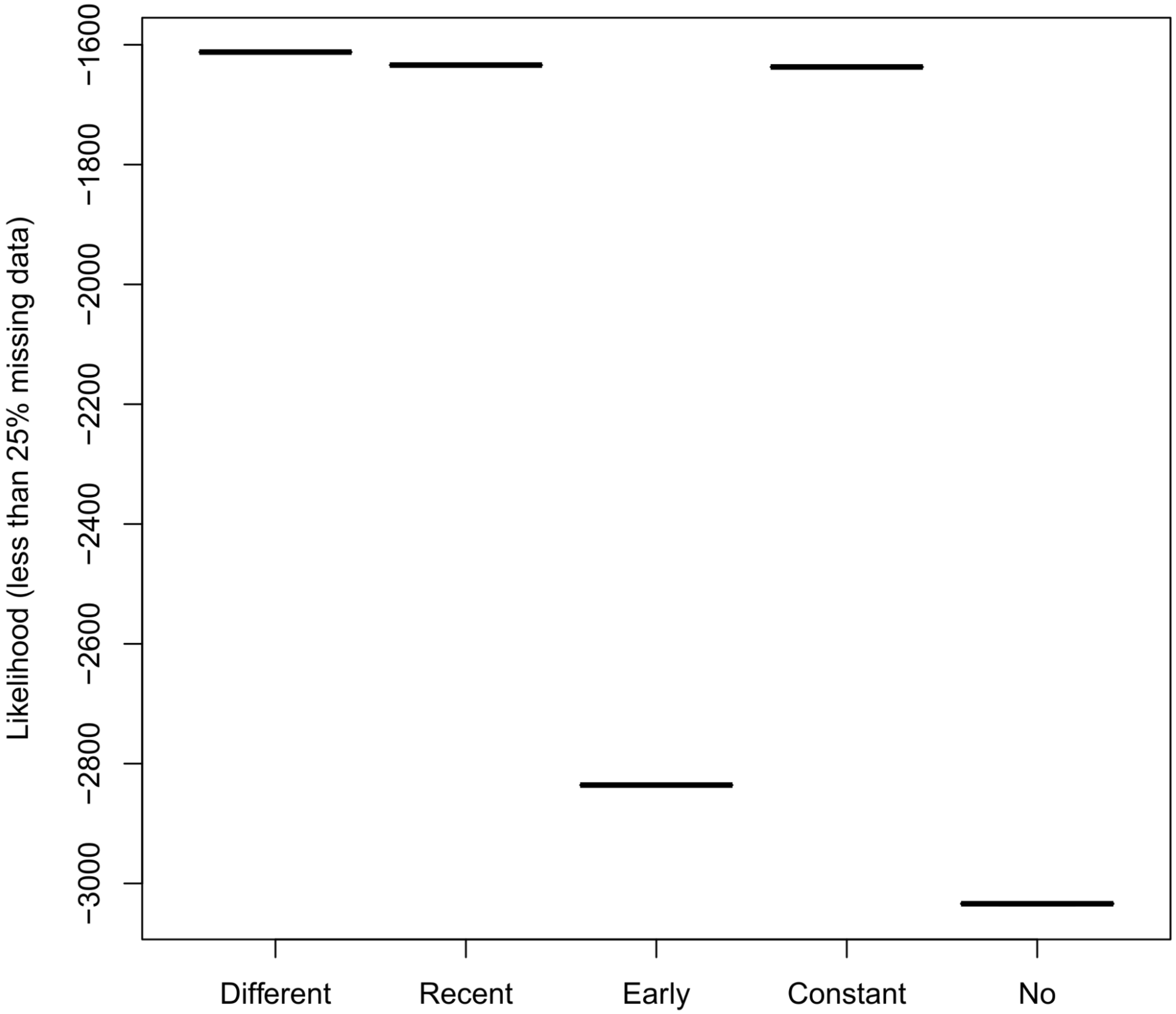


**Sup. Fig. 5.** Likelihood distribution for the *fastsimcoal* models with the SFS without the individuals with more than 25% missing data.

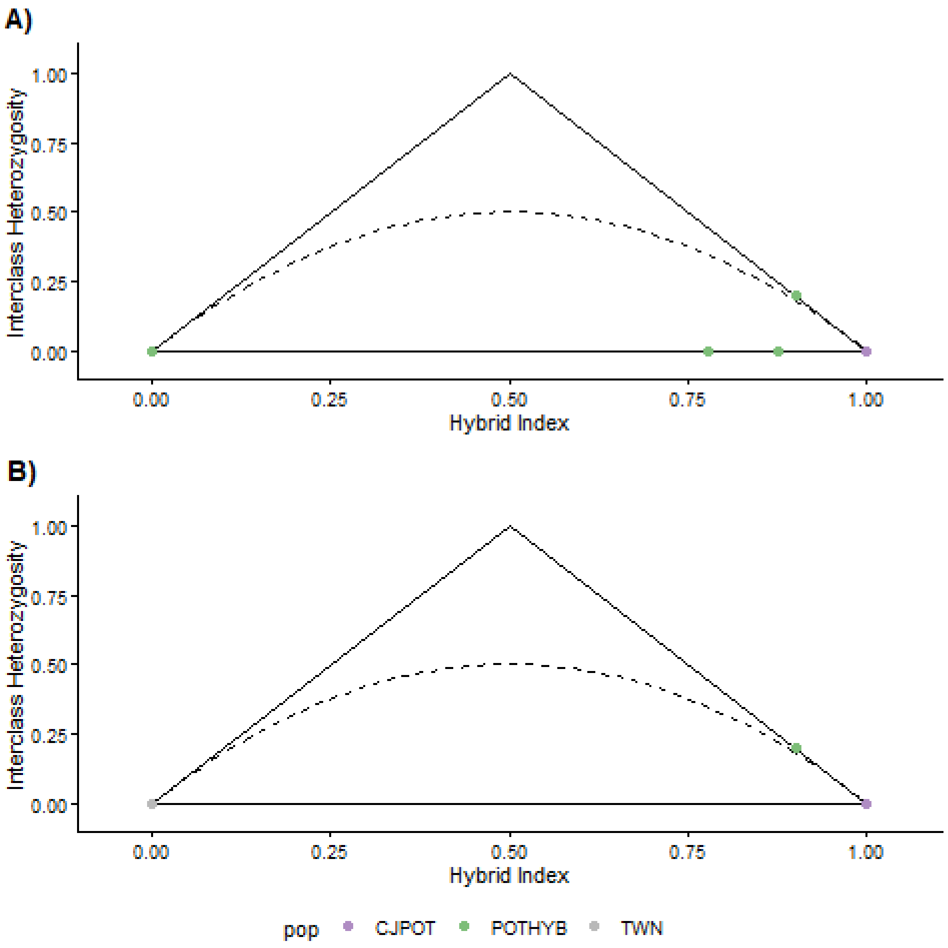


**Sup. Fig. 6.** A) Triangle plot for all the dataset. B) Triangle plot for the individuals with less than 25% missing data. “pop” = population. “CJPOT” = *C. japonensis* individuals according to ADMIXTURE. “POTHYB” = potential hybrids. “TWN” = *C. formosanus* individuals according to ADMIXTURE.
